# Structural Analysis of Genomic and Proteomic Signatures Reveal Dynamic Expression of Intrinsically Disordered Regions in Breast Cancer and Tissue

**DOI:** 10.1101/2023.02.23.529755

**Authors:** Nicole Zatorski, Yifei Sun, Abdulkadir Elmas, Christian Dallago, Timothy Karl, David Stein, Burkhard Rost, Kuan-Lin Huang, Martin Walsh, Avner Schlessinger

## Abstract

Structural features of proteins capture underlying information about protein evolution and function, which enhances the analysis of proteomic and transcriptomic data. Here we develop <u>S</u>tructural <u>A</u>nalysis of <u>G</u>ene and protein <u>E</u>xpression <u>S</u>ignatures (SAGES), a method that describes expression data using features calculated from sequence-based prediction methods and 3D structural models. We used SAGES, along with machine learning, to characterize tissues from healthy individuals and those with breast cancer. We analyzed gene expression data from 23 breast cancer patients and genetic mutation data from the COSMIC database as well as 17 breast tumor protein expression profiles. We identified prominent expression of intrinsically disordered regions in breast cancer proteins as well as relationships between drug perturbation signatures and breast cancer disease signatures. Our results suggest that SAGES is generally applicable to describe diverse biological phenomena including disease states and drug effects.

## 1. Introduction

With the advent of improved sequencing technology there has been a great emphasis placed on using proteomic and transcriptomic data to understand underlying disease etiologies and characteristics^1^. This has led to successful identification of biomarkers that have advanced precision medicine^2^ in fields such as oncology^3^. For example, single cell RNA sequencing on primary breast cancer tumors has explored the heterogeneity of gene expression in tumor tissue and the preponderance of immune cell response to disease^4^. Analysis of gene sets from RNA expression can be based on a comparison of the gene names found to be differentially expressed in a sample population as compared to the control population. The alternative, well-established method for analyzing sequencing data is through the use of a gene ontology (GO) enrichment of the differentially expressed genes, which provides a standardized vocabulary and relationship for genes^5^. This gives a slightly more nuanced view of the signature; in that it provides annotations about the function of proteins encoded by genes. In proteomic analysis, protein names or fragments of protein sequences found using mass spectroscopy are analyzed for their relative abundances in a sample^6^.

Despite the developments in transcriptomic and proteomic analysis tools, analysis of expression frequencies on a gene name or a GO annotation level does not capture functional similarities of genes based on their encoded protein structures. Structural features of proteins can robustly describe biological function. For example, we have shown that using one structural feature, i.e., the structural fold, family, and superfamily compositions of gene sets from transcriptomics data, captured similarities and differences among tissues and drug treated cell lines when combined with machine learning techniques^7^. Other showed that the use of structural information can detect off target effects of drugs when structurally similar proteins were targeted^8^. This holds true for structure-derived representations of proteins, such as secondary structure, solvent accessibility, and intrinsically disordered regions (IDRs)^9^. Accurate classifiers predicting structural features are available^9–13^, yielding 1D or string representations for input proteins. Moreover, the 3D structure prediction method AlphaFold2^14^ allows genome-scale 3D structure prediction, and representations from full 3D coordinates to distance and contact maps.

Here we tested whether representation of the gene or protein set with a variety of structural features can describe phenotypes. We first developed Structural Analysis of Gene and protein Expression Signatures (SAGES), a method of generating structural features from gene and protein sets. We applied SAGES, in combination with random forest models and recursive feature elimination, to GTEx^15^, a dataset of RNA expression data taken from human tissue samples. We also analyzed SAGES of breast cancer gene and protein expression data from patient cohorts, as well as breast cancer data from the COSMIC^16^ database of mutated and overexpressed genes. Furthermore, we applied SAGES drug perturbation signatures from the Connectivity map^17^ to investigate the similarity between existing breast cancer drugs and the breast cancer SAGES signature.

## 2. Results

### 2.1. Structural Features are Predictive of Normal Tissue Type

To test whether or not structural features can capture meaningful biological patterns, we evaluated the ability of SAGES to predict normal tissue from GTEx using only structural features and no gene name information. For each of the 30 tissue types in GTEx, we trained three random forest models using different input features, including structural features, gene names, and a combination of structural features with gene names. We measured average performance for all tissues following 10-fold cross validation (**Table 1**). Notably, accuracy and AUROC are 97% for all groups, including those trained on a combination of structural features and gene names. Interestingly, because performance of the model trained on genes was already high, a comparable performance with structural features alone that does not include any specific information on the genes is notable. For example, in the model trained on normal breast tissue SAGES the AUROC and F score were 0.949±0.012 and 0.951±0.011, respectively (**Figure 1**), while the gene name trained model obtained similar performance (0.951±0.012 and 0.953±0.011, respectively) (**Supplementary Table 2**). This demonstrates that, while gene names contain adequate information for recapitulating tissue type, remarkably, structural features alone also contain sufficient information for recapitulating biological identity of tissues. The breast tissue prediction model trained on both types of features preformed similarly to the structural features trained model with AUROC and F-score of 0.953±0.014 and 0.955±0.014 (**Figure 1**). This is unsurprising considering that it uses the same structural features in addition to the gene names. Predictor performance of normal tissue type using structural features compared to gene names alone demonstrates that the orthogonal information based on underlying protein characteristics is comparable to the information used in gene expression analysis.

**Figure 1.**
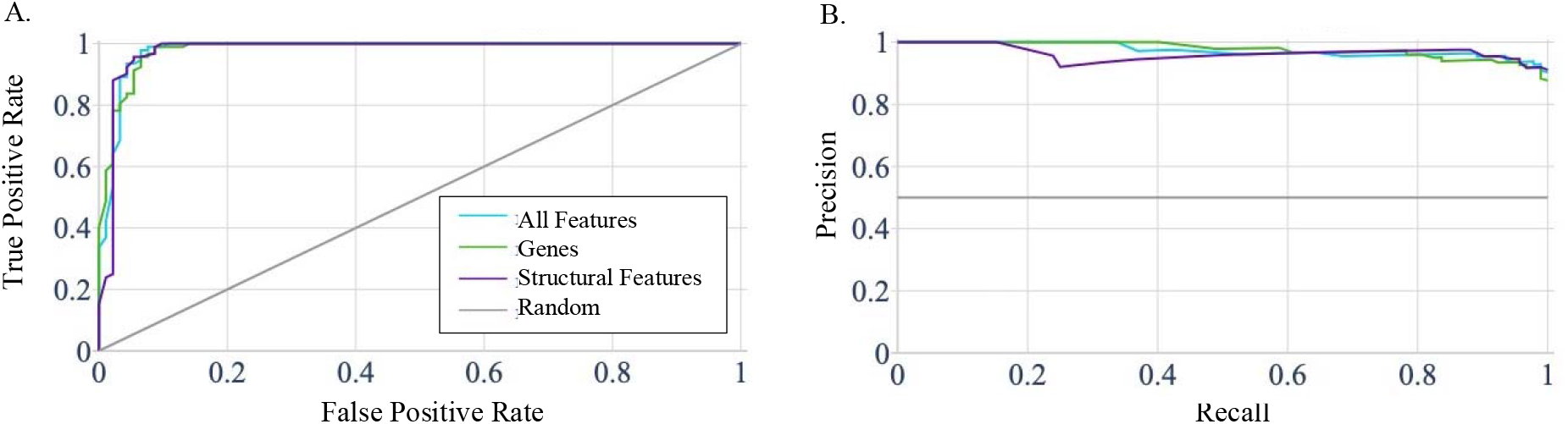
SAGES performance in predicting tissue from GTEx. Graphs depicting performance of the random forest predictor of normal breast tissue trained on structural features, gene names, and a combination of all features based on GTEx samples. The line labeled random represents performance of a model that has no skill and instead randomly selects a classification. (**A**) The Receiver Operating Characteristic Curve (ROC) for the breast tissue prediction model trained on different features, which all perform comparably. The AUROC for this model is 94.9% for training on structural features, 95.1% for training on gene names, and 95.3% for training on all features. (**B**) The precision recall curve for the breast tissue prediction model trained on different features. The precision and recall for each of the feature sets used to train the model are 92.8% and 97.5% for structural features, 92.6% and 98.0% for gene names, and 93.0% and 98.0% for all features combined.

**Table 1.** Performance of tissue type predictors with different input features. Averaged performance of random forest tissue type predictors on all thirty GTEx tissues following 10-fold cross validation. The Structural Features rows mark the performance obtained when the model was trained using only structural features. The gene name row denotes the performance of the model trained on one hot encoded gene name symbols. The structural features and gene names row represent the performance of the model trained on both structural features and gene names.

| Features Trained on | Tissue Type | Accuracy | AUROC | F score | Precision | Recall |
| --- | --- | --- | --- | --- | --- | --- |
| Structural Features and Gene Names | Average of all tissues | 0.972±0.061 | 0.972±0.061 | 0.974±0.054 | 0.971±0.071 | 0.98±0.049 |
| Gene Names | Average of all tissues | 0.973±0.057 | 0.973±0.057 | 0.975±0.054 | 0.972±0.068 | 0.979±0.051 |
| Structural Features | Average of all tissues | 0.967±0.063 | 0.967±0.063 | 0.97±0.054 | 0.965±0.077 | 0.979±0.049 |

### 2.2. Structural Features Reveal Characteristics of Normal Breast Tissue

To interrogate the contribution of different features to the model, we used recursive feature elimination (RFE)^18^ to analyze the normal breast tissue predictor trained on a combination of structural features and genes from GTEx samples (**Table 2**). Using all features as input to RFE allows us to explore the relative importance of genes and structural features when used in tandem. We observe that prominent genes are differentially expressed in normal breast tissue, including APOD, a component of HDL; FABP4, a fatty acid binding protein; SAA1, serum amyloid; and TIMP3, a metallopeptidase inhibitor. The identification of genes such as APOD and FABP4, which are known to be associated with breast tissue, increase our confidence in the result^19^. Other genes such as TIMP3 and SAA1 are known more for their breast cancer prognostic value^20,21^ and further support the robustness of the feature selection process.

**Table 2.**
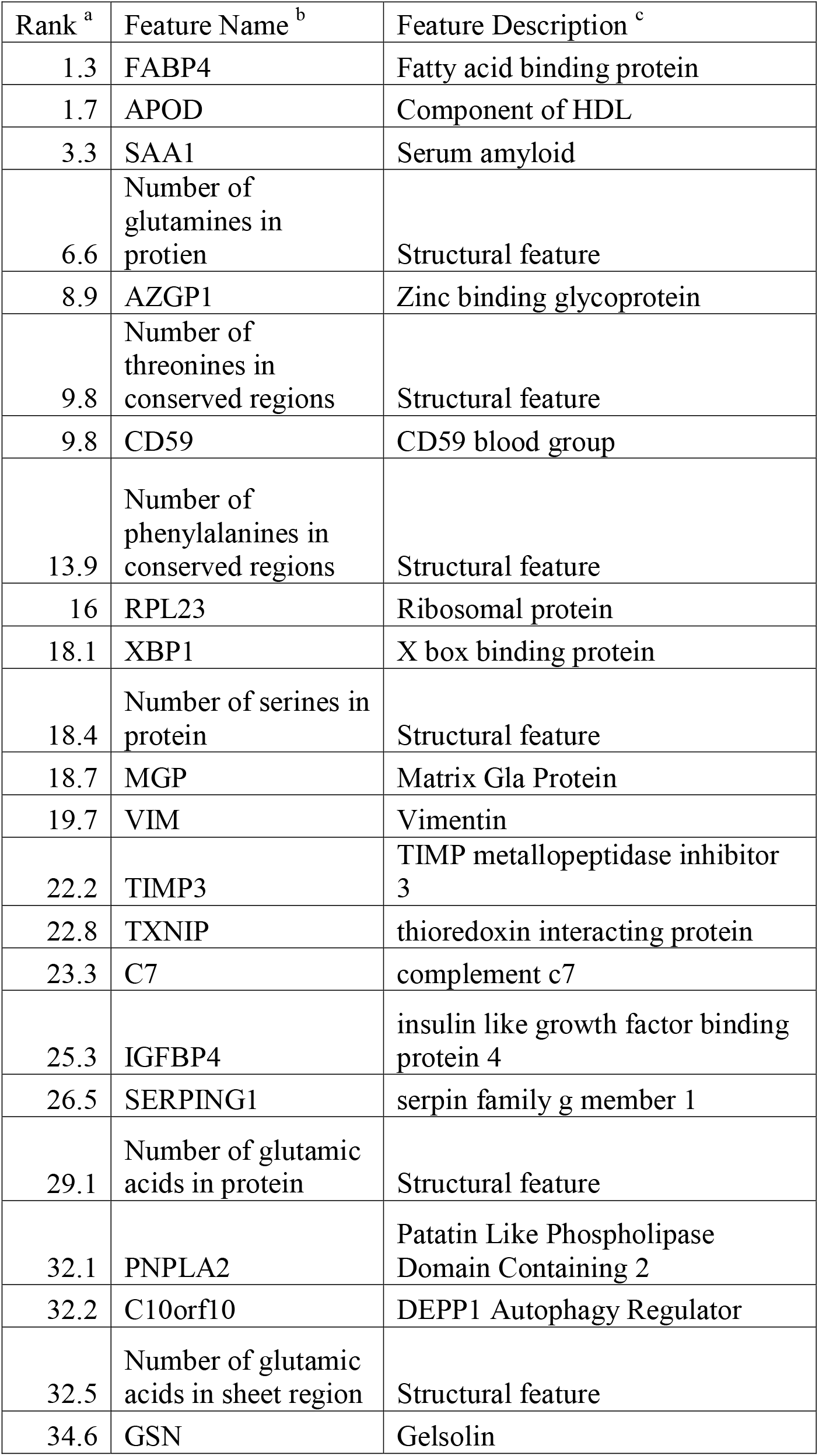
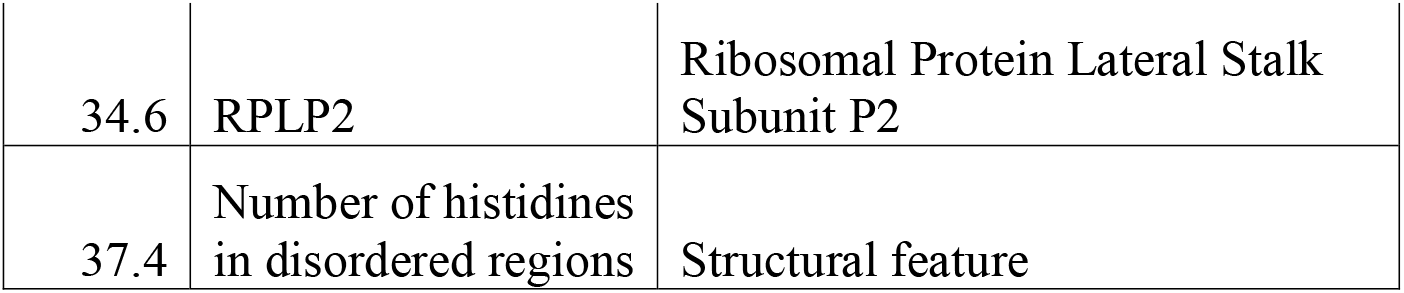
Ranking and description of features found most informative in the breast tissue predictor. **The twenty-five features that contributed most to the random forest model predicting normal breast tissue from structural features and gene names of GTEx samples.** ^a^ *Rank* marks the lowest value of rank corresponds to the feature that contributes most to the model as assigned using recursive feature elimination with 10 random seeds. ^b^ Feature Name represents contains the gene name symbol or structural feature name that corresponds with the rank in that same row. ^c^ Feature Description corresponds to additional information about the function or name of the feature in that row. Gene feature descriptors were summarized from the Human Gene Database Gene Cards version 5.10 [1].

Notably, features related to intrinsically disordered regions (IDRs) and conserved regions contributed greatly to the breast tissue predictor, even amongst all features, including gene names as well as other structural features (**Table 2**). These include the composition of glutamine, serine, and glutamate in the protein list; threonine and phenylalanine in conserved regions of proteins; glutamate residues in beta sheet regions; and histidine residues in the IDRs. In particular, this reveals that certain amino acids are differentially expressed in proteins that define breast tissue. For example, glutamine can potentially contribute to protein structural instability when found in IDRs^22^, and histidine, is known to be disorder promoting^23^. Taken together, this overall the importance of disorder related features is consistent with the enrichment of IDRs in signaling proteins, and particularly breast cancer, such as HER2^24^, SIPA1^25^, and BRCA1^26^.

### 2.3. Breast Cancer Structural Features Differ from those of Normal Breast Tissue

#### 2.3.1 Transcriptomic data reveals an overexpression of proteins with IDRs and intrinsically disordered binding regions: experimentally derived samples

Human breast tumor samples and normal breast tissue controls were collected from 23 individuals during their surgical treatment procedures and sequenced. We compared SAGES of these biopsied breast tumors to those of normal tissue samples from the same individual to identify changes in the type of protein features overexpressed in the diseased tissue. The SAGES analysis of all 23 patients, as well as the following subsets: HER2 negative patients, PR negative patients, ER negative patients, TCHP treated patients, and AC-T treated patients, all demonstrated statistically significantly different protein features from background. Interestingly, the most highly represented features in overexpressed genes in breast cancer patients are long IDRs and IDRs that interact with other proteins (intrinsically disordered binding regions) (**Figure 2A**). This trend is observed both in the agglomeration of all 23 samples and specifically in HER2 negative patients (**Figure 2B**). The important role of IDRs in tumorigenesis of a wide variety of cancers has previously been highlighted^27^. Many of the corresponding proteins with IDRs associated with cancer have been implicated in cell signaling pathways^28^. An examination of domains, folds, superfamilies, and families that are expressed in the breast cancer samples but not their corresponding normal breast tissues reveals cell signaling related substructures such as the PDZ domain^29^ and the tyrosine-protein kinase catalytic domain^30^. Additional features of note include zinc finger domains and immunoglobulin substructures, which are both implicated in breast cancer progression^31,32^. The immunoglobulin V-set domain is highly overexpressed in the set of breast cancer with PR negative disease and does not appear in HER2 or ER negative patients (**Figure 2B**). Interestingly, patients treated with TCHP had some features that were overexpressed compared to background (**Supplementary Figure 2A**), while those treated with AC-T only had features that were under expressed at a low magnitude compared to background (**Supplementary Figure 2B**). These trends provide motivation for further exploration of SAGES features in drug treatment.

**Figure 2.**
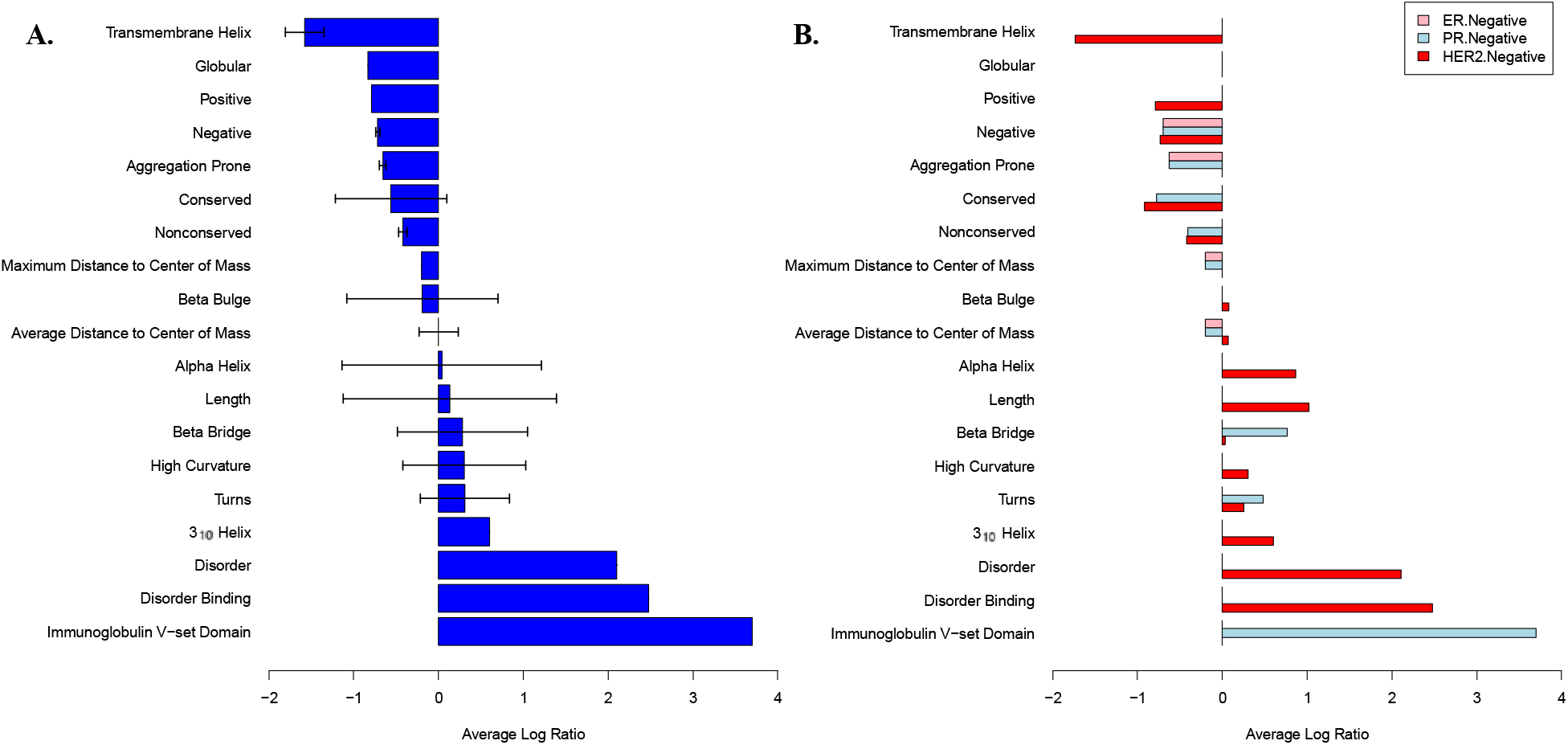
Structural features of normal and breast cancer tissue from newly generated human patient samples. Average log ratios of SAGES features that are statistically significantly different in breast cancer samples as compared to normal breast tissue. The log ratio of sample divided by background for each feature was averaged if the calculated p value for that feature was less than the Bonferroni adjusted significance level. Error bars represent the standard deviation of the values used to calculate the average. If there was only one instance of feature found to be statistically significant, the error bars were set to zero. Features that are not families, folds, superfamilies, domains, or frequency counts of a secondary structure consist of the number of amino acids within the protein that make up the secondary structure. (**A**) The average log ratio of SAGES features for gene expression from 23 breast cancer tumors compared to normal breast tissue from the same individuals. (**B**). The average log ratio of SAGES features for gene expression from 23 breast cancer patients separated by negative receptor status.

Unlike SAGES of most breast cancer patients that show features, which previously have been generalized to cancer appear in breast cancer, SAGES of triple negative disease (HER2, PR, and ER negative), Anastrozole treated disease, and GO analysis of all breast cancer samples did not reveal any statistically different gene ontology classifications between the cancerous and noncancerous breast tissue when also using a Bonferroni adjusted significance level.

#### 2.3.2 Transcriptomic data reveals an overexpression of proteins with IDR and intrinsically disordered binding regions: COSMIC database

SAGES captured breast cancer disease trends across datasets from multiple sources as seen in the analysis of breast cancer gene expression data from COSMIC^16^, an established cancer mutation database. Comparisons between mutated and unmutated but overexpressed breast cancer genes from COSMIC against normal breast tissue gene expression from GTEx demonstrated clear overexpression of IRDs and intrinsically disordered binding regions (**Figure 3**), in agreement with the analysis of the patient-derived breast cancer gene expression samples (**Figure 2**).

**Figure 3.**
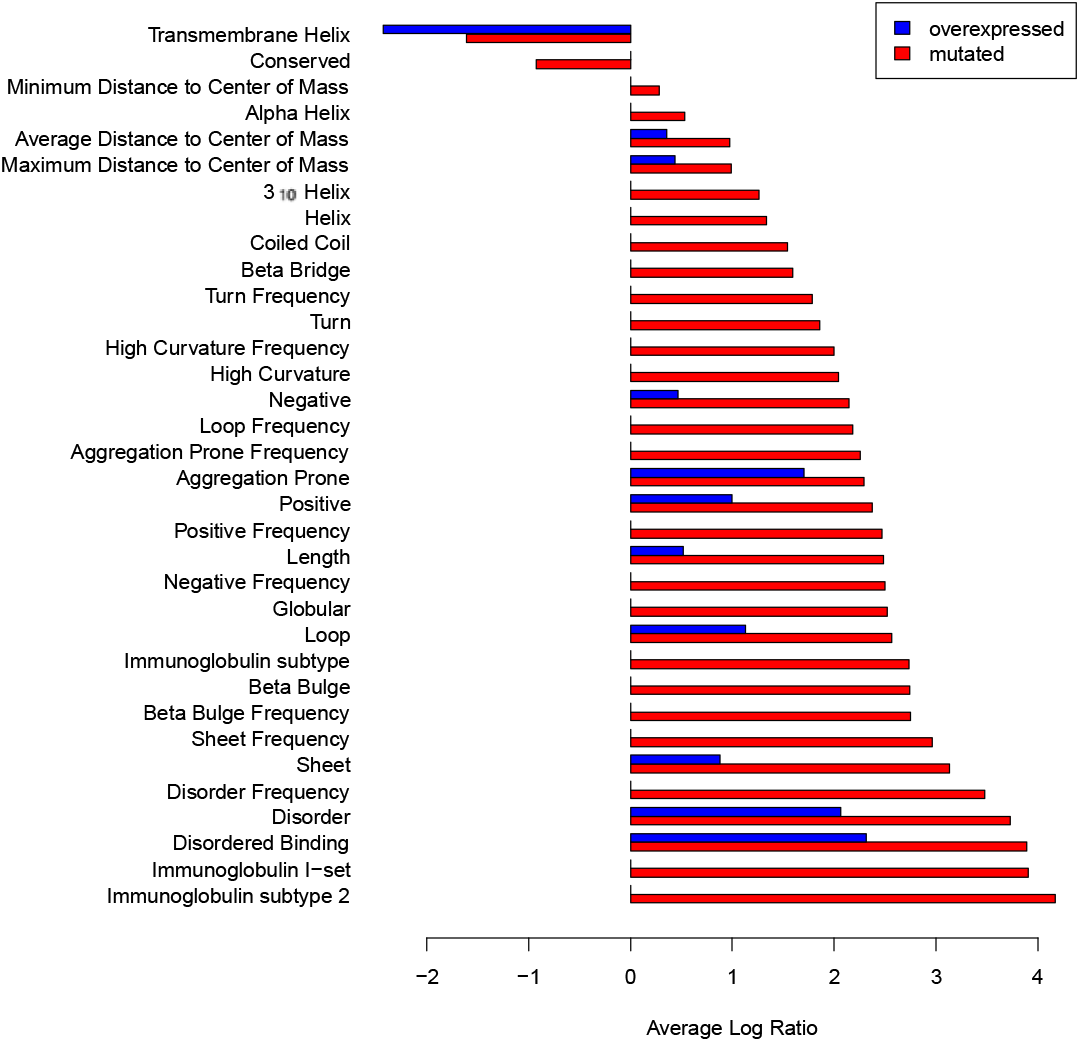
Structural features of normal and breast cancer tissue from the COSMIC database. The average log ratio of SAGES features for gene expression of mutated and unmutated but over expressed genes from COSMIC breast cancer tumors compared to normal breast tissue from GTEx. Features are displayed by size of average log ratio level for mutated genes, where positive values correspond to features with increased length or frequency in breast cancer compared to normal breast tissue. Due to the fact that the COSMIC data was based on a set of samples combined before analysis, there are no error bars.

Further, SAGES of the COSMIC data also provide insight into the difference between breast cancer genes that are mutated and those that are overexpressed but have unchanged amino acid sequences. Particularly, the most common mutations include: missense substitutions, nonsense substitutions, frameshift insertions, and frameshift deletions^16^. Both sets (i.e. mutated genes and overexpressed genes) over express proteins with long IDRs and intrinsically disordered binding regions, and the mutated genes are also enriched with Immunoglobulin Subtype domains as compared to normal breast tissue. Overall, the mutated genes are more different, in terms of number of statistically significantly different features and log ratio of expression, from normal background breast tissue than the unmutated genes. Interestingly, both mutated and overexpressed gene sets are enriched with transmembrane helices.

Addition of GO enrichment analysis further differentiates the roles associated with the mutated and unmutated but overexpressed breast cancer genes. Mutated breast cancer genes are associated with adhesion (such as 0007156, 0098742, 0031344), signaling (such as 0099177, 0050804, 0007267), cellular differentiation (such as 0000902, 0007420, 0048858), metabolism-including those of lipids, (such as 0019222, 0044237, 0044238), signaling related to immune response (such as 0002757, 0002429, 0002768), and regulation of cell death (0010941). Localization of these proteins primarily coincides with the plasma membrane (e.g. 0098590) and the extracellular region (e.g. 0005615). Notably, the mutated genes were implicated in processes related to adhesion. This is unsurprising due to the loss of cellular adhesion associated with most cancers^33^. Overexpressed breast cancer genes that have no amino acid sequence changes are associated with metabolism (such as 0008152, 0044238, 0006629), development (such as 0030154), signaling (such as 0023052, 0007165), regulation of cell death (0010942), and immune response (such as 0045087). Some of these proteins are associated with the extracellular space (0005615), however many others are associated with the endoplasmic reticulum and ribosomes (e.g. 0005788, 0005840). The localization to organelles related to protein synthesis is supported by the known increased metabolic activity of cancer cells^34^. Importantly, both of these RNA expression derived samples have processes related to signaling.

#### 2.3.3 Proteomic data reveals an overrepresentation of transmembrane proteins

Proteomic data, which can reveal differential protein abundance when compared to transcriptomic data^35^, provides a unique view of expressed breast cancer protein features. This supplements our understanding of breast cancer proteins, which are revealed through gene expression analysis. Proteomics data from 17 breast cancer tumors with normal tissues from the same patients was analyzed using SAGES^36^. The result showed the presence of proteins with transmembrane helix regions containing a larger number of amino acids on average and immunoglobulin domains (**Figure 4**). The proteomics data revealed different patterns in features from those calculated using gene expression signatures from the 23 different breast cancer patients sequenced for this study and the COSMIC gene expression data such as presence of the kinase regulatory SH3 domain^37^, with the exception of the immunoglobulin like domain. Interestingly, both domain types are often found in receptor tyrosine kinases (RTKs), which play a key role in signaling^38^. Notably, features from the proteomics breast cancer data set had shorter IDRs and binding regions that are intrinsically disordered. Because these partially intrinsically disordered proteins are often involved in cell signaling, such as RTKs, they are tightly regulated and have short half-life^39^. The difference in structural features found in the analysis of gene and protein expression of breast cancer therefore can be attributed to the transient nature of signaling proteins, which contain IDRs.

**Figure 4.**
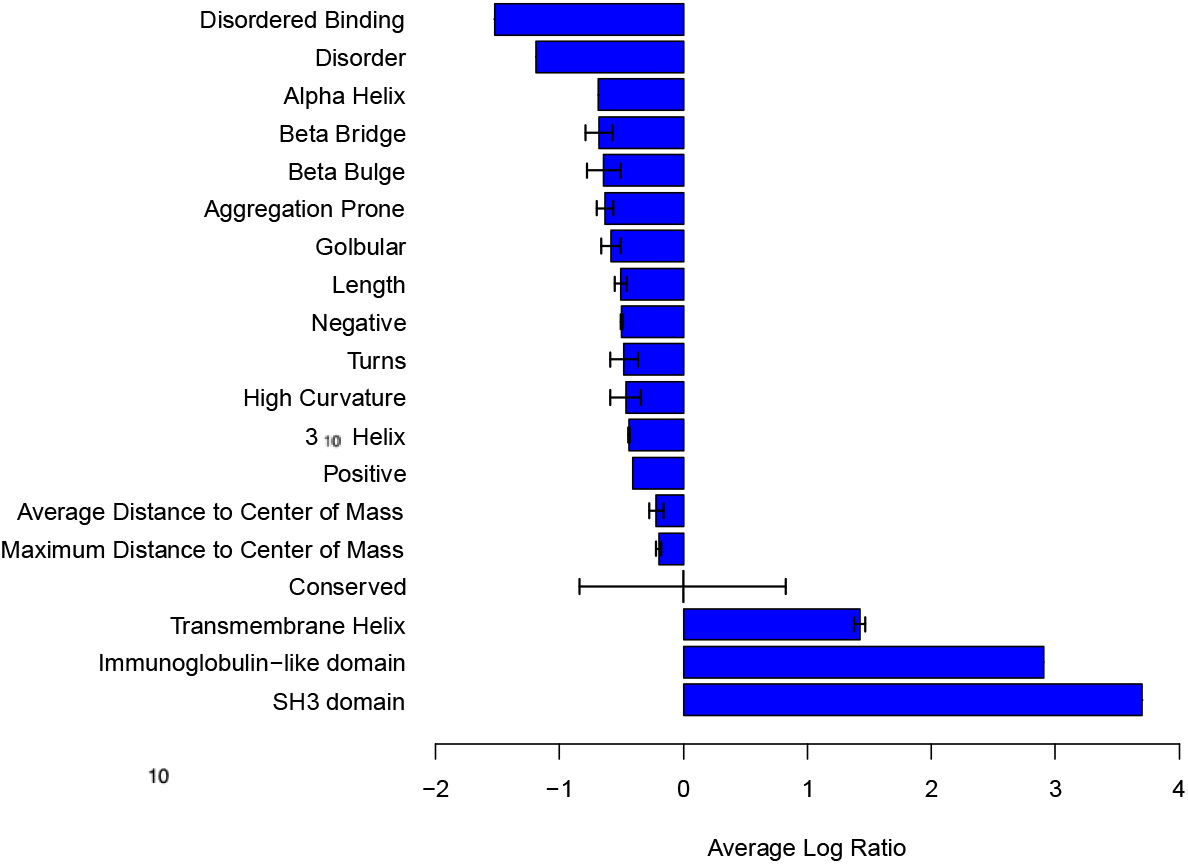
Structural features of primary breast tumors compared to matched normal tissues from a global proteomics dataset quantified using mass-spectrometry. The average log ratio of SAGES features for proteomics of 17 breast cancer tumors extracted from surgical patients compared to normal breast tissue from the same patients. Negative values correspond to features with decreased length or frequency in breast cancer compared to normal breast tissue.

This absence of transient signaling proteins in the proteomic sample is reflected in the GO analysis. On a biological process level, the proteins in breast cancer had functions related to metabolic processes (e.g. 0044281) and transcription (e.g. 0006351, 0097659). One of the areas where these proteins localize in the cell is the nucleus, specifically chromatin (0000785) and chromosomes (0005694). This aligns with the biological processes related to transcription. The proteins also localize to the extracellular space (0005615), which could be related to the increased presence of proteins with transmembrane helices, and the endoplasmic reticulum lumen (0031093). Notably absent from the overrepresented proteins found in the proteomic analysis of breast cancer are the GO terms related to any cellular signaling, as well as cellular adhesion. Taken together, our results suggest that combining analyzing using enrichments of both GO and structural features generated with SAGES supports the correlation between lack of signaling molecules and IDRs in the proteomic samples.

### 2.4. SAGES Captures Similarity Between Perturbation Signatures of Existing Breast Cancer Therapies and Disease

It has been proposed that gene signatures can be applied for drug repurposing^40^. In particular, the signature reversion principle (SRP) is based on the assumption that drug perturbation gene expression signature has a negative correlation with a disease gene expression signature^40^. To investigate the extension of the SRP, we applied SAGES to gene and protein expression changes induced by breast-cancer drugs and breast cancer. The number of statistically significantly different SAGES features between the drug perturbation and both genomic and proteomic breast cancer analyses were calculated and averaged for the two drug types. Comparisons between drug perturbation with proteomic and genomic breast cancer backgrounds show that breast cancer drugs produce a perturbation in cell lines with SAGES features that are more similar to those obtained from breast cancer than those obtained from other drugs (**Figure 5A**). With the proteomic background, the breast cancer drugs had a lower average number of SAGES features with statistically significant differences (314) than other drugs (321) (p=0.00070). With the genomic background, the breast cancer drugs had a lower average number of features with statistically significant differences (323) than other drugs (327) (p=0.050).

**Figure 5.**
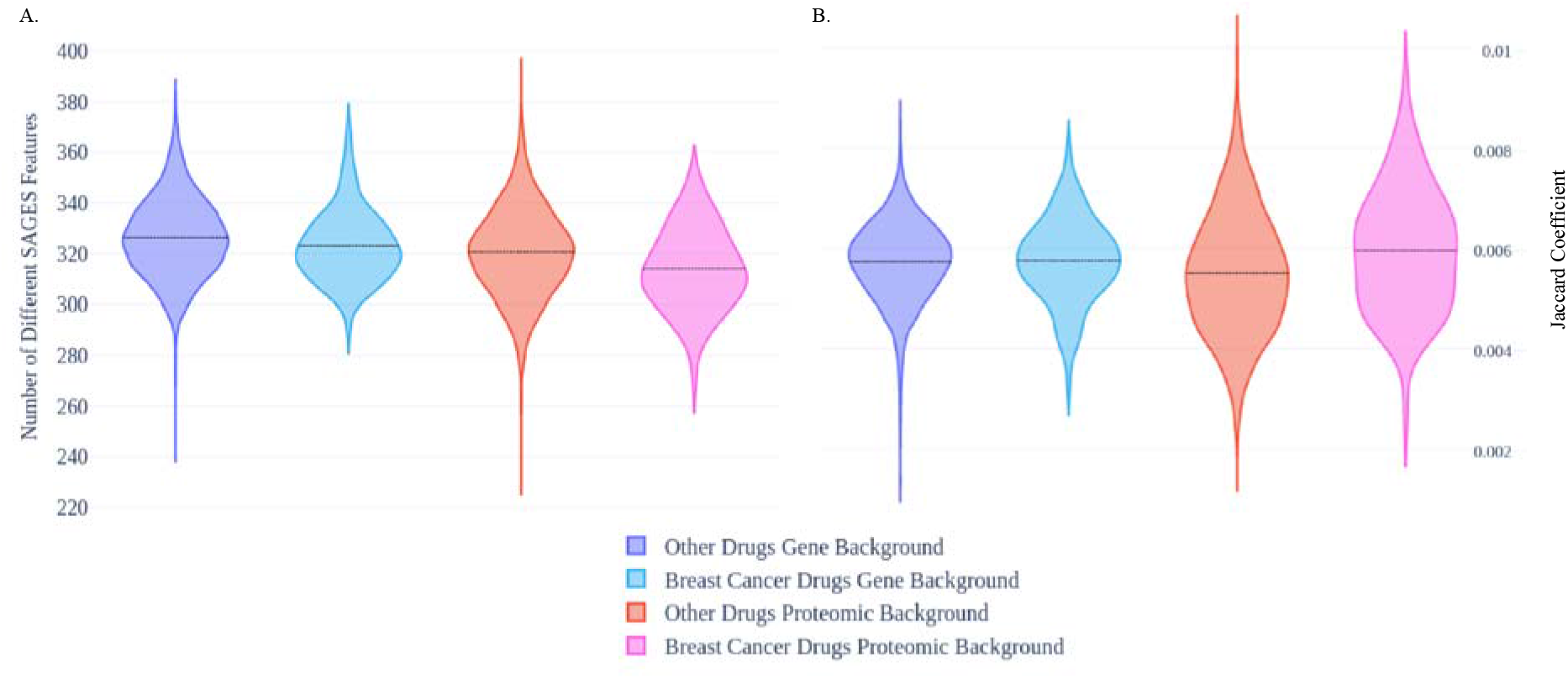
Perturbation signatures of existing breast cancer drugs and drugs for all other indications compared to breast cancer signatures. Difference between connectivity map breast cancer and other drug perturbation signatures compared to signatures derived from breast cancer COSMIC gene expression and experimental proteomics expression. (**A**). Counts of statistically significant different features between SAGES of breast cancer and drug perturbation (**B**) Jaccard coefficient representing similarity of drug perturbation signatures to gene and protein breast cancer signature backgrounds.

This increased similarity between drug perturbation and breast cancer is captured in a gene-protein level comparison between the perturbations and backgrounds but not in a gene-gene level comparison (**Figure 5B**). When the Jaccard coefficients of the breast cancer (0.0060) and other (0.0055) drug genes are compared to the proteomic breast cancer background, there is smore similarity to the background in the breast cancer drug genes (p=0.00102). However, this trend of SAGES features and proteins of breast cancer drug perturbations being more similar to the targeted disease is not recapitulated in the comparison of gene expression of drug perturbations and breast cancer for both drug groups. Breast cancer drugs have an average Jaccard coefficient of 0.00575 compared to other drugs with a Jaccard coefficient of 0.00572 (p=0.77). Because drug repurposing using perturbation signatures is classically done at a gene level^41^, the emergence of this association at the protein and structural level demonstrates the power of the orthogonal information provided with SAGES and raises a potential application of SAGES in the signature based repurposing sphere.

## 3. Discussion

Genomic and proteomic data contains valuable information about biological states. Currently, gene expression is assessed on a gene name similarity basis or through the use of enrichment of GO annotations in gene lists, however these existing methods do not capture underlying structure based functional information.

Application of features derived from different levels of protein structure can greatly enhance transcriptomic analysis. SAGES is distinct from existing methods in that it provides both sequence and 3D structure based orthogonal data which supplement transcriptomic biological insight (**Figure 6**). These detailed, structural level, descriptions of proteins encoded by the genes in the input protein data set provide orthogonal information to gene names and GO functional descriptions.

**Figure 6.**
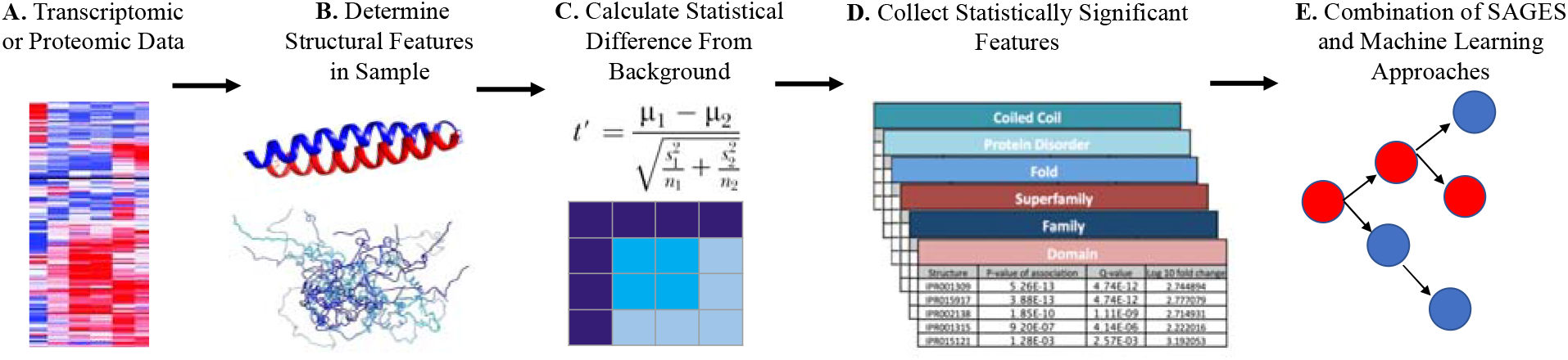
SAGES workflow. (**A**) SAGES takes as input a set of genes or proteins. (**B**) For each protein, SAGES determines structural features of protein regions from protein sequences and 3D models. (**C**) SAGES normalizes features by the magnitude of their corresponding protein’s expression level. (**D**) Sample structural features are then compared to a background using p values from a Fisher’s exact test or Welch’s t test. Bonferroni and false discovery rate corrected significance levels are provided based on the found feature size. (**E**) The output is a vectorized depiction of the structural features of a transcriptomics or proteomics sample that can be used as the input for biological analysis of samples and machine learning applications.

The analysis of normal tissue transcriptomic data from GTEx revealed the generalizability of structural features to all tissue types with the development of tissue type random forest predictors (Table 1). Use of features alone captured just as much information about biological state as the traditionally used gene names. Feature selection applied to the normal breast tissue samples further revealed that, interestingly, amino acid composition of IDRs was shown to play an informative role in breast tissue prediction (Table 2).

We applied SAGES to multiple breast cancer datasets to demonstrate how this method can be used to inform our understanding of breast tissue and tumor biology. SAGES, applied to samples excised from 23 women during their breast cancer treatment surgeries, revealed structural and functional differences between tumor and normal tissue. Notably, proteins with IDRs and intrinsically disordered binding regions are overexpressed in diseased breast cancer tissue (**Figure 2**). This was supported by SAGES analysis of breast cancer genes that are mutated and overexpressed in breast cancer according to COSMIC (**Figure 3**). GO analysis of these samples reveals that the proteins that are overexpressed in breast cancer and contain these IDR features are associated with cell signaling, metabolism, and immune response. Uniquely, mutated breast cancer proteins also were associated with cellular adhesion. Proteomic analysis of human breast cancer samples revealed a different feature landscape which was correlated with a loss of representation of proteins related to cell signaling (**Figure 4**). This is thought to be due to the transient nature of these cell signaling proteins.

SAGES was used to interrogate drug perturbation signatures from the Connectivity Map and the resulting analysis showed breast cancer drug perturbation signatures are more similar to breast cancer expression signatures on the feature and protein level than other drugs (**Figure 5**). This, along with SAGES analysis of notable features in breast cancer patients before receiving AC-T or TCHP treatment (Supplementary Figure 2), has interesting implications for signature reversion principle (SRP). This further demonstrates the capacity for structural features to capture both established and novel underlying biological function. The scope of SAGES is wide and we anticipate that it has the potential for a vast number of future applications.

## 4. Experimental Procedures

### 4.1. Structural Feature Generation

Transcriptomic signatures can be represented by the structural and functional features of the proteins transcribed in the gene set. These structural features can be derived from protein sequence or from three-dimensional, resolved structure. SAGES calculates sequence based features including length, frequency and amino acid composition from sequence predictions such as: IDRs predicted with IUPRED2a^42^; protein binding regions in intrinsically disordered peptides predicted with ANCHOR^43^; transmembrane helical regions predicted with TMHMM 2.0^44^; and globular, helical, positive amino acid containing regions, negative amino acid containing regions, coiled-coil, sheet, loop, conserved, and non-conserved regions predicted with PredictProtein^45^. SAGES also assigns SCOPe^46^ and UniProt^47^ folds, families, superfamilies, and domains using HHpred^48^, with optimized parameters determined in previous work^7^. SAGES applies the same amino acid frequency calculation to features derived from structural models generated by AlphaFold2^14^ (default parameters): 3_10_ helix, alpha helix, beta bridge, beta bulge, turns, and high curvature regions from the Dictionary of Protein Secondary Structure (DSSP)^49^; distances from the center of mass from biopython’s pdb parser^50^; and aggregation propensity from Aggrescan3d^51^. SAGES determines the number of amino acids (length), the number of separate instances of a feature type (frequency), and the number of each type of amino acid (composition) for secondary features. For all secondary features SAGES uses a cutoff of 50% predicted probability according to the various feature prediction tools listed above when determining amino acid content of a region. Unlike the other features, aggregation propensity is denoted with a range of values that can be negative or positive rather than a percentage of predicted probability so all amino acids with positive values are included in the analysis. Additionally, SAGES tabulates the total number of contacts as well as the minimum, maximum, and average amino acid distance from the protein’s center of mass. All features are listed in **Supplementary Table 1.**

SAGES normalizes the features using the expression level of the input genes or proteins which correspond to each feature. This weighted average normalization ensures feature frequency adequately reflects prominence within the sample. SAGES further normalizes the output feature values using the number of genes or proteins input in the sample to ensure consistency between different sized samples. Family, fold, superfamily, and domain were not normalized due to the underabundance of these features within each input set.

### 4.2. Statistical significance of features

The statistical tests included in the method compare the current sample to a background sample. The preloaded background consists of the entire human proteome from all GTEx V8 samples^15^. This background was replaced with a control background matched to each experimental sample in the breast cancer feature analysis. The frequencies of categorical variables are compared to the background using the scipy^52^ statistical package’s Fisher’s exact test to determine a p value. A two tailed, type two, t test was used to determine the t value for averages of lengths of secondary structures compared to background Eq. 1. Because variance between sample and background was not assumed to be equal, a Welch’s t-test was used^53^.

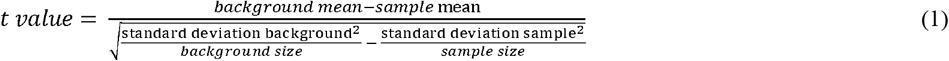

The scipy statistical package’s t.sf function with degrees of freedom equal to sample size minus two was used to determine the p value from the t value.

The p value cutoff was set at 0.05 and Bonferroni corrected^54^ according to the number of found features in that set of feature types. For families, folds, superfamilies, and domains, this means the number of each of those types of features found. For this correction, number of amino acids and secondary structure elements were grouped together and always summed to 219 leading to an adjusted p value of 0.000228. The p value cutoff for the averages was corrected by 13 resulting in an adjusted value of 0.00385. The statsmodels stats.multitest package’s false discovery rate correction^55^, was also calculated to provide an additional metric for determining significance for large samples. Base 2 log ratios of feature frequency in samples over background was also provided for categorical variables (**Figure 6**). The code for structural features can be found on the following GitHub: https://github.com/schlessinger-lab/structural_features

### 4.3. Prediction of Normal Tissues

Structural features were used to predict tissue type from RNA expression of normal tissues. Furthermore, the breast tissue model specifically was also used to highlight important structural characteristics of proteins that were highly predictive in this tissue type. The top 250 most highly expressed genes for each GTEx sample were extracted from GTEx V8^15^ along with their level of overexpression, which is defined as the log fold change greater than zero. As seen in previous work, this is a sufficiently sized number of genes for capturing underlying tissue specificity^7^. Structural features (section 2.1) for each sample were determined. There were 3,532 structural features and 5,376 gene name features for each sample. Sample features were min-max normalized (Eq. (2)) in preparation for the tissue type prediction where x_i_ equals the value of a feature for a sample and x equals the set of values for all samples corresponding to that feature.

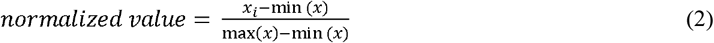

For each of the 30 tissue types, the samples were split into 90/10 training and test sets with equal number of positive and negative labels. The scikit learn version 0.24.1 random forest classifier was used to predict whether a sample could be classified as that type of tissue. This classifier fits 100 decision trees on data subsets and averages them into a meta predictor to improve performance. The following parameter values were used: criterion to measure split quality was gini impurity, maximum depth was none, minimum sample split was two, minimum samples leaves was one, minimum weight fraction leaf was zero, maximum features was the square root of the number of features, maximum leaf nodes was none, minimum impurity decrease was zero, bootstrap samples were used to build the trees, out of bag samples were not used to estimate the generalization score, number of parallel jobs was none, verbosity was none, warm start was set to false and just fit a new forest every call, class weights were none, complexity parameter used for pruning was zero, and maximum samples was none. Following 10-fold cross validation with random seed 0-9, performance was measured using: the area under the receiver operating characteristic curve (AUROC), accuracy Eq. (3) where *y_i_* is the value of the i^th^ sample and 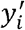 is the corresponding predicted value, precision Eq. (4) where TP is true positives and FP is false positives, recall Eq. (5) where FN is false negatives, and F-score Eq. (6). Standard deviations for all metrics were computed using python.statistics stdev.

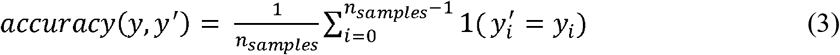

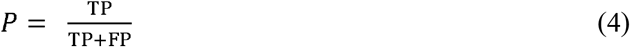

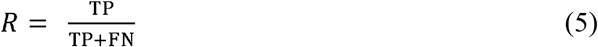

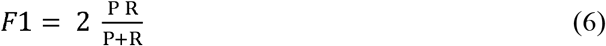

Additionally, the most predictive features for normal breast tissue were determined using sklearn recursive feature elimination (RFE) with random forest as the estimator^18^. Weights were assigned to all features and then the less important features were eliminated recursively until a single feature remained. This generated a ranking of features based on contribution to the model. RFE was conducted 10 times with random seeds 0-9 and rankings were summed. The 25 features with the lowest score, and therefore best rank are reported in Table 2.

### 4.4. Breast Cancer Datasets: Generation

Breast tumor tissue and normal control breast tissue was excised from 23 breast cancer patients during surgery. Samples were then prepared and sequenced according to standard protocol^56^. The women enrolled had the following demographics: 13 were post-menopausal, 4 were Estrogen Receptor (ER) negative, 16 were Human Epidermal Growth Factor Receptor (HER2) negative, 5 were Progesterone Receptor (PR) negative, 16 had invasive ductal carcinoma, and 2 had internal mammary chain involvement. Five women received treatment prior to sample collection. One woman was administered Taxotere, Carboplatin, Herceptin, Perjeta (TCHP). Three women were administered Adriamycin, cyclophosphamide, Taxol (AC-T). One woman was administered Anastrozole. Global proteomics data from primary breast tumors and matched normal tissues were generated through mass-spectrometry (MS) (PMID: 33212010)^36^ and pre-processed as previously described (PMID: 34552204). Data pertaining to mutated and overexpressed genes in human breast cancer was also sourced from the Catalogue of Somatic Mutations In Cancer (COSMIC)^16^. This was divided into two sets, one containing only the most commonly mutated genes in breast cancer and the other containing only unmutated, overexpressed genes in breast cancer. For weighting purposes, the number of times a gene was mutated compared to the total number of times that gene was expressed served as the weight for the first set and the number of times the gene was overexpressed was used as the weight for the second set.

### 4.5. Breast Cancer Datasets: Analysis

The 250 most highly expressed genes were extracted from each genomic and proteomic sample and SAGES was applied. Each patient breast tumor sample had a matching normal breast tissue control from the same patient, which was used to generate the background for the statistical analysis described as part of the SAGES method. The COSMIC samples were compared to SAGES of the top 250 genes from all GTEx normal breast tissue samples. For each sample, if a feature was statistically significant (p value less than the Bonferroni corrected significance level of 0.0011 determined by dividing 0.05 by the number of non-amino acid specific features), the log ratio was included in the averages visualized in Figure 3. The standard deviation was determined for all features that came from more than one sample. Due to the large number of amino acid related features, these were excluded from the visualization. Folds, superfamilies, and domains were present only in the breast cancer samples and not in the background were excluded from this analysis.

Gene Ontology (GO) analysis was also performed using the top 250 most overexpressed genes for each dataset with the panther classification system and the Bonferroni adjusted significance level^57^. The following sample-background pairs were compared using this approach: breast cancer transcriptomic samples-normal breast transcriptomic samples, breast cancer proteomic samples-normal breast proteomic samples, breast cancer proteomic samples-breast cancer transcriptomic samples, normal breast tissue proteomic samples-normal breast tissue transcriptomic samples, COSMIC breast cancer samples with and without mutations-GTEx normal breast tissue.

### 4.6. Breast Cancer Drug Perturbation

Perturbation signatures of cell lines treated with drugs from the Connectivity Map^1^ were compared to various breast cancer backgrounds to investigate how drug induced gene expression relates to disease signature. Unweighted SAGES for all overexpressed genes for each experimental sample were calculated and compared to unweighted SAGES of all proteins from the proteomics samples with log ratio expression greater than one and to unweighted SAGES of all genes from the COSMIC mutated breast cancer gene dataset. Unweighted SAGES were used to ensure that all overexpressed proteins contributed equally to the analysis. For each sample, the number of significantly different features from background were counted. The significance level of 0.05 was selected. Multiple hypothesis testing was not employed because the aim was ultimately to assess similarity to background rather than difference from background and using this technique would increase the type II error. The samples were divided into breast cancer treatment drugs (doxorubicin, fulvestrant, letrozole, megestrol, methotrexate, paclitaxel, raloxifene, tamoxifen, and vinblastine) according to a list published by the National Cancer Institute, and all other drugs in the Connectivity Map database. The average number of statistically significantly different features for each group were calculated and a two-sided, type 2, student t test was used to determine the p value.

Gene expression of the perturbation samples and the expression signatures used to calculate the SAGES of the two backgrounds were also directly compared. The Jaccard coefficient, which is the intersection of the genes overexpressed in both sets over the union of the genes overexpressed in both sets, was used to determine signature similarity. Samples were divided into breast cancer and other drugs and compared with a two-sided, type 2, student t test. Additionally, the average Jaccard coefficient for both groups was determined.

## Acknowledgments

We would like to thank Michael Bernhofer and Nolan Caile for their assistance. This work is supported by the National Institutes of Health grant F30 HL160179 to N.Z.

## Contributions

N.Z. developed and used SAGES to analyze the datasets. N.Z. and A.S. wrote the manuscript. Y.S. and M.W. generated the breast cancer gene expression data. A.E. and K.L.H. provided the processed proteomic breast cancer data. N.Z., C.D., M.B., T.K., B.R., and D.S. contributed to feature generation.

## Declarations

The authors declare no competing interests

