## Supplemental Figures for "Structural Analysis of Genomic and Proteomic Signatures Reveal Dynamic Expression of Intrinsically Disordered Regions in Breast Cancer and Tissue"

**Supplementary Figure 1**

See separate PDF

**Supplementary Figure 1.** Graphs depicting performance of the random forest predictor of normal tissues trained on structural features, gene names, and a combination of all features based on GTEx samples. The line labeled random represents performance of a model that has no skill and instead randomly selects a classification. The Receiver Operating Characteristic Curve (right) and Precision Recall Curve (left) for each tissue’s random seed 0 predictor performance.

**Supplementary Figure 2**

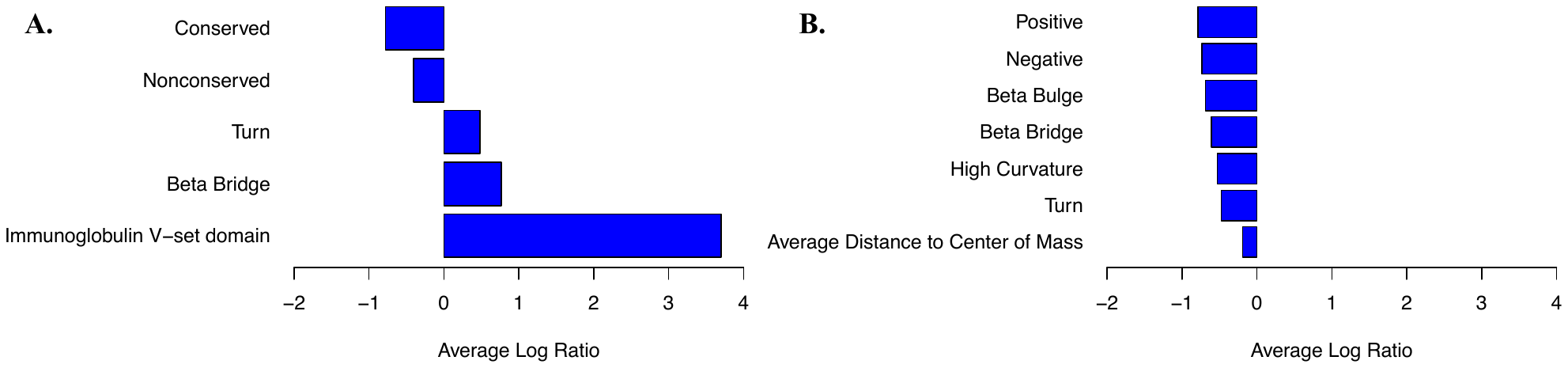

**Supplementary Figure 2. SAGES from gene expression data of breast cancer patients before starting breast cancer chemotherapy regimens.** A. Statistically significantly different structural features in breast cancer samples in the TCHP treated population. B. Statistically significantly different structural features in breast cancer samples in the AC-T treated population

Supplementary Table 1.

Domains

SCOPe Families

SCOPe Superfamilies

SCOPe Folds

Length of protein

Negative amino acid region lengths

Positive amino acid region lengths

Anchor (disorder binding) region lengths

Coil region lengths

Conserved region lengths

Disordered region lengths

Globular region lengths

Helix region lengths

Loop region lengths

Nonconserved region lengths

Sheet region lengths

Transmembrane helix region lengths

High curvature region lengths

Beta bulge region lengths

Turn region lengths

Beta Bridge region lengths

3_10_ helix region lengths

Alpha helix region lengths

Aggregation Prone region lengths

Minimum distance to the center of mass

Maximum distance to the center of mass

Average distance to the center of mass

Number of amino acids in anchor regions: A

Number of amino acids in anchor regions: C

Number of amino acids in anchor regions: D

Number of amino acids in anchor regions: E

Number of amino acids in anchor regions: F

Number of amino acids in anchor regions: G

Number of amino acids in anchor regions: H

Number of amino acids in anchor regions: I

Number of amino acids in anchor regions: K

Number of amino acids in anchor regions: L

Number of amino acids in anchor regions: M

Number of amino acids in anchor regions: N

Number of amino acids in anchor regions: P

Number of amino acids in anchor regions: Q

Number of amino acids in anchor regions: R

Number of amino acids in anchor regions: S

Number of amino acids in anchor regions: T

Number of amino acids in anchor regions: V

Number of amino acids in anchor regions: W

Number of amino acids in anchor regions: Y

Number of amino acids in coil regions: A

Number of amino acids in coil regions: C

Number of amino acids in coil regions: D

Number of amino acids in coil regions: E

Number of amino acids in coil regions: F

Number of amino acids in coil regions: G

Number of amino acids in coil regions: H

Number of amino acids in coil regions: I

Number of amino acids in coil regions: K

Number of amino acids in coil regions: L

Number of amino acids in coil regions: M

Number of amino acids in coil regions: N

Number of amino acids in coil regions: P

Number of amino acids in coil regions: Q

Number of amino acids in coil regions: R

Number of amino acids in coil regions: S

Number of amino acids in coil regions: T

Number of amino acids in coil regions: V

Number of amino acids in coil regions: W

Number of amino acids in coil regions: Y

Number of amino acids in conserved regions: A

Number of amino acids in conserved regions: C

Number of amino acids in conserved regions: D

Number of amino acids in conserved regions: E

Number of amino acids in conserved regions: F

Number of amino acids in conserved regions: G

Number of amino acids in conserved regions: H

Number of amino acids in conserved regions: I

Number of amino acids in conserved regions: K

Number of amino acids in conserved regions: L

Number of amino acids in conserved regions: M

Number of amino acids in conserved regions: N

Number of amino acids in conserved regions: P

Number of amino acids in conserved regions: Q

Number of amino acids in conserved regions: R

Number of amino acids in conserved regions: S

Number of amino acids in conserved regions: T

Number of amino acids in conserved regions: V

Number of amino acids in conserved regions: W

Number of amino acids in conserved regions: Y

Number of amino acids in disordered regions: A

Number of amino acids in disordered regions: C

Number of amino acids in disordered regions: D

Number of amino acids in disordered regions: E

Number of amino acids in disordered regions: F

Number of amino acids in disordered regions: G

Number of amino acids in disordered regions: H

Number of amino acids in disordered regions: I

Number of amino acids in disordered regions: K

Number of amino acids in disordered regions: L

Number of amino acids in disordered regions: M

Number of amino acids in disordered regions: N

Number of amino acids in disordered regions: P

Number of amino acids in disordered regions: Q

Number of amino acids in disordered regions: R

Number of amino acids in disordered regions: S

Number of amino acids in disordered regions: T

Number of amino acids in disordered regions: V

Number of amino acids in disordered regions: W

Number of amino acids in disordered regions: Y

Number of amino acids in globular regions: A

Number of amino acids in globular regions: C

Number of amino acids in globular regions: D

Number of amino acids in globular regions: E

Number of amino acids in globular regions: F

Number of amino acids in globular regions: G

Number of amino acids in globular regions: H

Number of amino acids in globular regions: I

Number of amino acids in globular regions: K

Number of amino acids in globular regions: L

Number of amino acids in globular regions: M

Number of amino acids in globular regions: N

Number of amino acids in globular regions: P

Number of amino acids in globular regions: Q

Number of amino acids in globular regions: R

Number of amino acids in globular regions: S

Number of amino acids in globular regions: T

Number of amino acids in globular regions: V

Number of amino acids in globular regions: W

Number of amino acids in globular regions: Y

Number of amino acids in helix regions: A

Number of amino acids in helix regions: C

Number of amino acids in helix regions: D

Number of amino acids in helix regions: E

Number of amino acids in helix regions: F

Number of amino acids in helix regions: G

Number of amino acids in helix regions: H

Number of amino acids in helix regions: I

Number of amino acids in helix regions: K

Number of amino acids in helix regions: L

Number of amino acids in helix regions: M

Number of amino acids in helix regions: N

Number of amino acids in helix regions: P

Number of amino acids in helix regions: Q

Number of amino acids in helix regions: R

Number of amino acids in helix regions: S

Number of amino acids in helix regions: T

Number of amino acids in helix regions: V

Number of amino acids in helix regions: W

Number of amino acids in helix regions: Y

Number of amino acids in loop regions: A

Number of amino acids in loop regions: C

Number of amino acids in loop regions: D

Number of amino acids in loop regions: E

Number of amino acids in loop regions: F

Number of amino acids in loop regions: G

Number of amino acids in loop regions: H

Number of amino acids in loop regions: I

Number of amino acids in loop regions: K

Number of amino acids in loop regions: L

Number of amino acids in loop regions: M

Number of amino acids in loop regions: N

Number of amino acids in loop regions: P

Number of amino acids in loop regions: Q

Number of amino acids in loop regions: R

Number of amino acids in loop regions: S

Number of amino acids in loop regions: T

Number of amino acids in loop regions: V

Number of amino acids in loop regions: W

Number of amino acids in loop regions: Y

Number of amino acids in non-conserved regions: A

Number of amino acids in non-conserved regions: C

Number of amino acids in non-conserved regions: D

Number of amino acids in non-conserved regions: E

Number of amino acids in non-conserved regions: F

Number of amino acids in non-conserved regions: G

Number of amino acids in non-conserved regions: H

Number of amino acids in non-conserved regions: I

Number of amino acids in non-conserved regions: K

Number of amino acids in non-conserved regions: L

Number of amino acids in non-conserved regions: M

Number of amino acids in non-conserved regions: N

Number of amino acids in non-conserved regions: P

Number of amino acids in non-conserved regions: Q

Number of amino acids in non-conserved regions: R

Number of amino acids in non-conserved regions: S

Number of amino acids in non-conserved regions: T

Number of amino acids in non-conserved regions: V

Number of amino acids in non-conserved regions: W

Number of amino acids in non-conserved regions: Y

Number of amino acids in protein: A

Number of amino acids in protein: C

Number of amino acids in protein: D

Number of amino acids in protein: E

Number of amino acids in protein: F

Number of amino acids in protein: G

Number of amino acids in protein: H

Number of amino acids in protein: I

Number of amino acids in protein: K

Number of amino acids in protein: L

Number of amino acids in protein: M

Number of amino acids in protein: N

Number of amino acids in protein: P

Number of amino acids in protein: Q

Number of amino acids in protein: R

Number of amino acids in protein: S

Number of amino acids in protein: T

Number of amino acids in protein: V

Number of amino acids in protein: W

Number of amino acids in protein: Y

Number of amino acids in sheet regions: A

Number of amino acids in sheet regions: C

Number of amino acids in sheet regions: D

Number of amino acids in sheet regions: E

Number of amino acids in sheet regions: F

Number of amino acids in sheet regions: G

Number of amino acids in sheet regions: H

Number of amino acids in sheet regions: I

Number of amino acids in sheet regions: K

Number of amino acids in sheet regions: L

Number of amino acids in sheet regions: M

Number of amino acids in sheet regions: N

Number of amino acids in sheet regions: P

Number of amino acids in sheet regions: Q

Number of amino acids in sheet regions: R

Number of amino acids in sheet regions: S

Number of amino acids in sheet regions: T

Number of amino acids in sheet regions: V

Number of amino acids in sheet regions: W

Number of amino acids in sheet regions: Y

Number of anchor regions

Number of coils

Number of conserved regions

Number of disordered regions

Number of globular regions

Number of helices

Number of loops

Number of regions with negative charge

Number of regions with negative charge with length >=30

Number of non-conserved regions

Number of regions with positive charge

Number of regions with positive charge with length >=30

Number of sheets

Number of transmembrane helices

Y/n anchor regions

Y/n disordered regions

Y/n globular regions

Y/n tmh regions

Number of high curvature regions

Number of amino acids in high curvature regions: A

Number of amino acids in high curvature regions: R

Number of amino acids in high curvature regions: N

Number of amino acids in high curvature regions: D

Number of amino acids in high curvature regions: C

Number of amino acids in high curvature regions: E

Number of amino acids in high curvature regions: Q

Number of amino acids in high curvature regions: G

Number of amino acids in high curvature regions: H

Number of amino acids in high curvature regions: I

Number of amino acids in high curvature regions: L

Number of amino acids in high curvature regions: K

Number of amino acids in high curvature regions: M

Number of amino acids in high curvature regions: F

Number of amino acids in high curvature regions: P

Number of amino acids in high curvature regions: S

Number of amino acids in high curvature regions: T

Number of amino acids in high curvature regions: W

Number of amino acids in high curvature regions: Y

Number of amino acids in high curvature regions: V

Number of beta bulge regions

Number of amino acids in beta bulge regions: A

Number of amino acids in beta bulge regions: R

Number of amino acids in beta bulge regions: N

Number of amino acids in beta bulge regions: D

Number of amino acids in beta bulge regions: C

Number of amino acids in beta bulge regions: E

Number of amino acids in beta bulge regions: Q

Number of amino acids in beta bulge regions: G

Number of amino acids in beta bulge regions: H

Number of amino acids in beta bulge regions: I

Number of amino acids in beta bulge regions: L

Number of amino acids in beta bulge regions: K

Number of amino acids in beta bulge regions: M

Number of amino acids in beta bulge regions: F

Number of amino acids in beta bulge regions: P

Number of amino acids in beta bulge regions: S

Number of amino acids in beta bulge regions: T

Number of amino acids in beta bulge regions: W

Number of amino acids in beta bulge regions: Y

Number of amino acids in beta bulge regions: V

Number of turn regions

Number of amino acids in turn regions: A

Number of amino acids in turn regions: R

Number of amino acids in turn regions: N

Number of amino acids in turn regions: D

Number of amino acids in turn regions: C

Number of amino acids in turn regions: E

Number of amino acids in turn regions: Q

Number of amino acids in turn regions: G

Number of amino acids in turn regions: H

Number of amino acids in turn regions: I

Number of amino acids in turn regions: L

Number of amino acids in turn regions: K

Number of amino acids in turn regions: M

Number of amino acids in turn regions: F

Number of amino acids in turn regions: P

Number of amino acids in turn regions: S

Number of amino acids in turn regions: T

Number of amino acids in turn regions: W

Number of amino acids in turn regions: Y

Number of amino acids in turn regions: V

Number of beta bridge regions

Number of amino acids in beta bridge regions: A

Number of amino acids in beta bridge regions: R

Number of amino acids in beta bridge regions: N

Number of amino acids in beta bridge regions: D

Number of amino acids in beta bridge regions: C

Number of amino acids in beta bridge regions: E

Number of amino acids in beta bridge regions: Q

Number of amino acids in beta bridge regions: G

Number of amino acids in beta bridge regions: H

Number of amino acids in beta bridge regions: I

Number of amino acids in beta bridge regions: L

Number of amino acids in beta bridge regions: K

Number of amino acids in beta bridge regions: M

Number of amino acids in beta bridge regions: F

Number of amino acids in beta bridge regions: P

Number of amino acids in beta bridge regions: S

Number of amino acids in beta bridge regions: T

Number of amino acids in beta bridge regions: W

Number of amino acids in beta bridge regions: Y

Number of amino acids in beta bridge regions: V

Number of 3_10_ helix regions

Number of amino acids in 3_10_ helix regions: A

Number of amino acids in 3_10_ helix regions: R

Number of amino acids in 3_10_ helix regions: N

Number of amino acids in 3_10_ helix regions: D

Number of amino acids in 3_10_ helix regions: C

Number of amino acids in 3_10_ helix regions: E

Number of amino acids in 3_10_ helix regions: Q

Number of amino acids in 3_10_ helix regions: G

Number of amino acids in 3_10_ helix regions: H

Number of amino acids in 3_10_ helix regions: I

Number of amino acids in 3_10_ helix regions: L

Number of amino acids in 3_10_ helix regions: K

Number of amino acids in 3_10_ helix regions: M

Number of amino acids in 3_10_ helix regions: F

Number of amino acids in 3_10_ helix regions: P

Number of amino acids in 3_10_ helix regions: S

Number of amino acids in 3_10_ helix regions: T

Number of amino acids in 3_10_ helix regions: W

Number of amino acids in 3_10_ helix regions: Y

Number of amino acids in 3_10_ helix regions: V

Number of alpha helix regions

Number of amino acids in alpha helix regions: A

Number of amino acids in alpha helix regions: R

Number of amino acids in alpha helix regions: N

Number of amino acids in alpha helix regions: D

Number of amino acids in alpha helix regions: C

Number of amino acids in alpha helix regions: E

Number of amino acids in alpha helix regions: Q

Number of amino acids in alpha helix regions: G

Number of amino acids in alpha helix regions: H

Number of amino acids in alpha helix regions: I

Number of amino acids in alpha helix regions: L

Number of amino acids in alpha helix regions: K

Number of amino acids in alpha helix regions: M

Number of amino acids in alpha helix regions: F

Number of amino acids in alpha helix regions: P

Number of amino acids in alpha helix regions: S

Number of amino acids in alpha helix regions: T

Number of amino acids in alpha helix regions: W

Number of amino acids in alpha helix regions: Y

Number of amino acids in alpha helix regions: V

Number of aggregation prone regions

Number of amino acids in aggregation prone regions: A

Number of amino acids in aggregation prone regions: R

Number of amino acids in aggregation prone regions: N

Number of amino acids in aggregation prone regions: D

Number of amino acids in aggregation prone regions: C

Number of amino acids in aggregation prone regions: E

Number of amino acids in aggregation prone regions: Q

Number of amino acids in aggregation prone regions: G

Number of amino acids in aggregation prone regions: H

Number of amino acids in aggregation prone regions: I

Number of amino acids in aggregation prone regions: L

Number of amino acids in aggregation prone regions: K

Number of amino acids in aggregation prone regions: M

Number of amino acids in aggregation prone regions: F

Number of amino acids in aggregation prone regions: P

Number of amino acids in aggregation prone regions: S

Number of amino acids in aggregation prone regions: T

Number of amino acids in aggregation prone regions: W

Number of amino acids in aggregation prone regions: Y

Number of amino acids in aggregation prone regions: V

Number of Contacts

Supplementary Table 1. List of all SAGES features

Supplementary Table 2.

| **Features Trained** | **Tissue Type** | **Accuracy** | **AUROC** | **F score** | **Precision** | **Recall** |
| --- | --- | --- | --- | --- | --- | --- |
| Structural Features and Gene Names | Adipose Tissue | 0.982±0.005 | 0.982±0.005 | 0.982±0.005 | 0.974±0.007 | 0.99±0.007 |
| Gene Names | Adipose Tissue | 0.985±0.004 | 0.985±0.004 | 0.986±0.003 | 0.977±0.007 | 0.994±0.003 |
| Structural Features | Adipose Tissue | 0.982±0.005 | 0.982±0.005 | 0.982±0.005 | 0.975±0.007 | 0.989±0.007 |
| Structural Features and Gene Names | Adrenal Gland | 0.999±0.003 | 0.999±0.003 | 0.999±0.003 | 1.0±0.0 | 0.998±0.006 |
| Gene Names | Adrenal Gland | 1.0±0.0 | 1.0±0.0 | 1.0±0.0 | 1.0±0.0 | 1.0±0.0 |
| Structural Features | Adrenal Gland | 1.0±0.0 | 1.0±0.0 | 1.0±0.0 | 1.0±0.0 | 1.0±0.0 |
| Structural Features and Gene Names | Bladder | 0.89±0.057 | 0.89±0.057 | 0.892±0.058 | 0.877±0.086 | 0.92±0.103 |
| Gene Names | Bladder | 0.89±0.099 | 0.89±0.099 | 0.894±0.094 | 0.877±0.117 | 0.92±0.103 |
| Structural Features | Bladder | 0.88±0.079 | 0.88±0.079 | 0.887±0.063 | 0.876±0.124 | 0.92±0.103 |
| Structural Features and Gene Names | Blood | 1.0±0.0 | 1.0±0.0 | 1.0±0.0 | 1.0±0.0 | 1.0±0.0 |
| Gene Names | Blood | 1.0±0.0 | 1.0±0.0 | 1.0±0.0 | 1.0±0.0 | 1.0±0.0 |
| Structural Features | Blood | 1.0±0.0 | 1.0±0.0 | 1.0±0.0 | 1.0±0.0 | 1.0±0.0 |
| Structural Features and Gene Names | Blood Vessel | 0.985±0.005 | 0.985±0.005 | 0.985±0.005 | 0.993±0.006 | 0.976±0.009 |
| Gene Names | Blood Vessel | 0.987±0.005 | 0.987±0.005 | 0.987±0.005 | 0.995±0.005 | 0.979±0.007 |
| Structural Features | Blood Vessel | 0.981±0.005 | 0.981±0.005 | 0.981±0.005 | 0.99±0.006 | 0.973±0.009 |
| Structural Features and Gene Names | Brain | 1.0±0.0 | 1.0±0.0 | 1.0±0.0 | 1.0±0.0 | 0.999±0.001 |
| Gene Names | Brain | 1.0±0.0 | 1.0±0.0 | 1.0±0.0 | 1.0±0.0 | 0.999±0.001 |
| Structural Features | Brain | 1.0±0.0 | 1.0±0.0 | 1.0±0.0 | 1.0±0.0 | 0.999±0.001 |
| Structural Features and Gene Names | Breast | 0.953±0.014 | 0.953±0.014 | 0.955±0.014 | 0.93±0.02 | 0.98±0.01 |
| Gene Names | Breast | 0.951±0.012 | 0.951±0.012 | 0.953±0.011 | 0.926±0.018 | 0.98±0.009 |
| Structural Features | Breast | 0.949±0.012 | 0.949±0.012 | 0.951±0.011 | 0.928±0.018 | 0.975±0.01 |
| Structural Features and Gene Names | Cervix | 0.838±0.084 | 0.838±0.084 | 0.848±0.076 | 0.823±0.131 | 0.9±0.129 |
| Gene Names | Cervix | 0.85±0.099 | 0.85±0.099 | 0.857±0.095 | 0.832±0.123 | 0.9±0.129 |
| Structural Features | Cervix | 0.838±0.084 | 0.838±0.084 | 0.851±0.077 | 0.803±0.116 | 0.925±0.121 |
| Structural Features and Gene Names | Colon | 0.983±0.005 | 0.983±0.005 | 0.983±0.005 | 0.983±0.01 | 0.983±0.007 |
| Gene Names | Colon | 0.979±0.008 | 0.979±0.008 | 0.979±0.008 | 0.973±0.015 | 0.985±0.011 |
| Structural Features | Colon | 0.98±0.008 | 0.98±0.008 | 0.98±0.008 | 0.98±0.011 | 0.98±0.01 |
| Structural Features and Gene Names | Esophagus | 0.98±0.004 | 0.98±0.004 | 0.98±0.004 | 0.979±0.007 | 0.981±0.008 |
| Gene Names | Esophagus | 0.98±0.004 | 0.98±0.004 | 0.98±0.004 | 0.98±0.005 | 0.981±0.008 |
| Structural Features | Esophagus | 0.977±0.005 | 0.977±0.005 | 0.977±0.005 | 0.975±0.009 | 0.978±0.008 |
| Structural Features and Gene Names | Fallopian Tube | 0.825±0.206 | 0.825±0.206 | 0.857±0.175 | 0.8±0.219 | 0.95±0.158 |
| Gene Names | Fallopian Tube | 0.85±0.175 | 0.85±0.175 | 0.87±0.164 | 0.817±0.2 | 0.95±0.158 |
| Structural Features | Fallopian Tube | 0.75±0.118 | 0.75±0.118 | 0.793±0.091 | 0.717±0.158 | 0.95±0.158 |
| Structural Features and Gene Names | Heart | 0.999±0.001 | 0.999±0.001 | 0.999±0.001 | 1.0±0.0 | 0.998±0.003 |
| Gene Names | Heart | 0.999±0.002 | 0.999±0.002 | 0.999±0.002 | 1.0±0.0 | 0.997±0.003 |
| Structural Features | Heart | 0.999±0.001 | 0.999±0.001 | 0.999±0.001 | 1.0±0.0 | 0.998±0.003 |
| Structural Features and Gene Names | Kidney | 0.968±0.049 | 0.968±0.049 | 0.969±0.045 | 0.964±0.071 | 0.979±0.048 |
| Gene Names | Kidney | 0.989±0.024 | 0.989±0.024 | 0.989±0.026 | 1.0±0.0 | 0.979±0.048 |
| Structural Features | Kidney | 0.954±0.051 | 0.954±0.051 | 0.956±0.047 | 0.945±0.08 | 0.971±0.05 |
| Structural Features and Gene Names | Liver | 1.0±0.0 | 1.0±0.0 | 1.0±0.0 | 1.0±0.0 | 1.0±0.0 |
| Gene Names | Liver | 0.986±0.03 | 0.986±0.03 | 0.985±0.032 | 1.0±0.0 | 0.971±0.06 |
| Structural Features | Liver | 1.0±0.0 | 1.0±0.0 | 1.0±0.0 | 1.0±0.0 | 1.0±0.0 |
| Structural Features and Gene Names | Lung | 0.999±0.002 | 0.999±0.002 | 0.999±0.002 | 1.0±0.0 | 0.997±0.004 |
| Gene Names | Lung | 0.998±0.002 | 0.998±0.002 | 0.998±0.002 | 0.999±0.003 | 0.997±0.004 |
| Structural Features | Lung | 0.998±0.002 | 0.998±0.002 | 0.998±0.002 | 1.0±0.0 | 0.996±0.005 |
| Structural Features and Gene Names | Muscle | 1.0±0.0 | 1.0±0.0 | 1.0±0.0 | 1.0±0.0 | 1.0±0.0 |
| Gene Names | Muscle | 1.0±0.0 | 1.0±0.0 | 1.0±0.0 | 1.0±0.0 | 1.0±0.0 |
| Structural Features | Muscle | 1.0±0.001 | 1.0±0.001 | 1.0±0.001 | 1.0±0.0 | 0.999±0.002 |
| Structural Features and Gene Names | Nerve | 0.996±0.003 | 0.996±0.003 | 0.996±0.003 | 1.0±0.0 | 0.992±0.005 |
| Gene Names | Nerve | 0.999±0.002 | 0.999±0.002 | 0.999±0.002 | 1.0±0.0 | 0.998±0.004 |
| Structural Features | Nerve | 0.992±0.004 | 0.992±0.004 | 0.991±0.004 | 0.999±0.003 | 0.984±0.008 |
| Structural Features and Gene Names | Ovary | 0.972±0.023 | 0.972±0.023 | 0.971±0.025 | 0.992±0.013 | 0.953±0.049 |
| Gene Names | Ovary | 0.969±0.022 | 0.969±0.022 | 0.968±0.024 | 0.992±0.013 | 0.947±0.048 |
| Structural Features | Ovary | 0.981±0.021 | 0.981±0.021 | 0.98±0.022 | 0.992±0.013 | 0.969±0.044 |
| Structural Features and Gene Names | Pancreas | 1.0±0.0 | 1.0±0.0 | 1.0±0.0 | 1.0±0.0 | 1.0±0.0 |
| Gene Names | Pancreas | 1.0±0.0 | 1.0±0.0 | 1.0±0.0 | 1.0±0.0 | 1.0±0.0 |
| Structural Features | Pancreas | 1.0±0.0 | 1.0±0.0 | 1.0±0.0 | 1.0±0.0 | 1.0±0.0 |
| Structural Features and Gene Names | Pituitary | 0.996±0.005 | 0.996±0.005 | 0.996±0.005 | 1.0±0.0 | 0.991±0.009 |
| Gene Names | Pituitary | 0.997±0.004 | 0.997±0.004 | 0.997±0.004 | 1.0±0.0 | 0.995±0.008 |
| Structural Features | Pituitary | 0.996±0.006 | 0.996±0.006 | 0.996±0.006 | 0.998±0.006 | 0.993±0.009 |
| Structural Features and Gene Names | Prostate | 0.972±0.023 | 0.972±0.023 | 0.972±0.024 | 0.978±0.029 | 0.967±0.035 |
| Gene Names | Prostate | 0.971±0.017 | 0.971±0.017 | 0.971±0.018 | 0.976±0.022 | 0.967±0.031 |
| Structural Features | Prostate | 0.958±0.027 | 0.958±0.027 | 0.958±0.028 | 0.965±0.025 | 0.951±0.041 |
| Structural Features and Gene Names | Salivary Gland | 0.989±0.012 | 0.989±0.012 | 0.99±0.012 | 0.985±0.02 | 0.994±0.013 |
| Gene Names | Salivary Gland | 0.989±0.01 | 0.989±0.01 | 0.99±0.01 | 0.983±0.02 | 0.997±0.01 |
| Structural Features | Salivary Gland | 0.988±0.014 | 0.988±0.014 | 0.988±0.014 | 0.985±0.021 | 0.991±0.015 |
| Structural Features and Gene Names | Skin | 0.998±0.001 | 0.998±0.001 | 0.998±0.001 | 1.0±0.001 | 0.996±0.002 |
| Gene Names | Skin | 0.998±0.002 | 0.998±0.002 | 0.998±0.002 | 1.0±0.001 | 0.995±0.003 |
| Structural Features | Skin | 0.997±0.001 | 0.997±0.001 | 0.997±0.001 | 1.0±0.001 | 0.995±0.003 |
| Structural Features and Gene Names | Small Intestine | 0.966±0.019 | 0.966±0.019 | 0.966±0.018 | 0.962±0.031 | 0.971±0.019 |
| Gene Names | Small Intestine | 0.961±0.021 | 0.961±0.021 | 0.961±0.02 | 0.957±0.04 | 0.966±0.018 |
| Structural Features | Small Intestine | 0.968±0.011 | 0.968±0.011 | 0.969±0.011 | 0.967±0.026 | 0.971±0.023 |
| Structural Features and Gene Names | Spleen | 0.993±0.007 | 0.993±0.007 | 0.993±0.007 | 0.988±0.014 | 0.998±0.006 |
| Gene Names | Spleen | 0.995±0.005 | 0.995±0.005 | 0.995±0.005 | 0.992±0.01 | 0.998±0.006 |
| Structural Features | Spleen | 0.991±0.011 | 0.991±0.011 | 0.991±0.011 | 0.984±0.022 | 0.998±0.006 |
| Structural Features and Gene Names | Stomach | 0.975±0.01 | 0.975±0.01 | 0.975±0.011 | 0.982±0.015 | 0.968±0.017 |
| Gene Names | Stomach | 0.972±0.014 | 0.972±0.014 | 0.972±0.014 | 0.97±0.023 | 0.974±0.019 |
| Structural Features | Stomach | 0.969±0.011 | 0.969±0.011 | 0.969±0.011 | 0.977±0.022 | 0.961±0.019 |
| Structural Features and Gene Names | Testis | 0.999±0.003 | 0.999±0.003 | 0.999±0.003 | 1.0±0.0 | 0.997±0.006 |
| Gene Names | Testis | 0.997±0.004 | 0.997±0.004 | 0.997±0.004 | 1.0±0.0 | 0.995±0.007 |
| Structural Features | Testis | 0.997±0.006 | 0.997±0.006 | 0.997±0.006 | 1.0±0.0 | 0.993±0.012 |
| Structural Features and Gene Names | Thyroid | 0.997±0.005 | 0.997±0.005 | 0.997±0.005 | 1.0±0.0 | 0.993±0.01 |
| Gene Names | Thyroid | 0.996±0.004 | 0.996±0.004 | 0.996±0.004 | 1.0±0.0 | 0.992±0.008 |
| Structural Features | Thyroid | 0.998±0.003 | 0.998±0.003 | 0.998±0.003 | 1.0±0.0 | 0.996±0.005 |
| Structural Features and Gene Names | Uterus | 0.981±0.019 | 0.981±0.019 | 0.981±0.019 | 0.98±0.023 | 0.983±0.024 |
| Gene Names | Uterus | 0.978±0.02 | 0.978±0.02 | 0.978±0.019 | 0.974±0.033 | 0.983±0.018 |
| Structural Features | Uterus | 0.971±0.028 | 0.971±0.028 | 0.971±0.028 | 0.96±0.031 | 0.983±0.034 |
| Structural Features and Gene Names | Vagina | 0.941±0.024 | 0.941±0.024 | 0.941±0.024 | 0.931±0.033 | 0.953±0.034 |
| Gene Names | Vagina | 0.939±0.029 | 0.939±0.029 | 0.939±0.029 | 0.936±0.036 | 0.944±0.035 |
| Structural Features | Vagina | 0.933±0.031 | 0.933±0.031 | 0.933±0.033 | 0.931±0.041 | 0.938±0.055 |

**Supplementary Table 2.** Performance of random forest tissue type predictors on all thirty GTEx tissues following 10-fold cross validation. The Structural Features rows mark the performance obtained when the model was trained using only structural features. The gene name row denotes the performance of the model trained on one hot encoded gene name symbols. The structural features and gene names row represent the performance of the model trained on both structural features and gene names.
